# Comparison study of sixteen differential abundance testing methods using two large Parkinson disease gut microbiome datasets

**DOI:** 10.1101/2021.02.24.432717

**Authors:** Zachary D. Wallen

## Abstract

**Background:** When studying the relationship between the microbiome and a disease, a common question asked is what individual microbes are differentially abundant between a disease and healthy state. Numerous differential abundance (DA) testing methods exist and range from standard statistical tests to methods specifically designed for microbiome data. Comparison studies of DA testing methods have been performed, but none were performed on microbiome datasets collected for the study of real, complex disease. Due to this, we performed DA testing of microbial genera using 16 DA methods in two large, uniformly collected gut microbiome datasets on Parkinson disease (PD), and compared their results.

**Results:** Pairwise concordances between methods ranged from 46%-99% similarity. Average pairwise concordance per dataset was 76%, and dropped to 62% when taking replication of signals across datasets into account. Certain methods consistently resulted in above average concordances (e.g. Kruskal-Wallis, ALDEx2, GLM with centered-log-ratio transform), while others consistently resulted in lower than average concordances (e.g. edgeR, fitZIG). Overall, ∼80% of genera tested were detected as differentially abundant by at least one method in each dataset. Requiring associations to replicate across datasets reduced significant signals by almost half. Further requirement of signals to be replicated by the majority of methods (≥8) yielded 19 associations. Only one genus (*Agathobacter*) was replicated by all methods. Use of hierarchical clustering revealed three groups of DA signatures that were (1) replicated by the majority of methods and included genera previously associated with PD, (2) replicated by few or no methods, and (3) replicated by a subset of methods and included rarer genera, all enriched in PD.

**Conclusions:** Differential abundance tests yielded varied results. Using one method on one dataset may find true associations, but may also detect non-reproducible signals, adding to inconsistency in the literature. To help lower false positives, one might analyze data with two or more DA methods to gauge concordance, and use a built-in replication dataset to show reproducibility. This study corroborated previously reported microorganism associations in PD, and revealed a potential new group of microorganisms whose abundance is significantly elevated in PD, and might be worth pursuing in future investigations.

## Background

Microbiome research has gained immense traction in recent years driven primarily by technological advances in sequencing and exponential increase in computational resources and tools. The availability of these new tools and technologies have solidified a place for microbiome research in many fields of research including the biomedical research community where a large portion of the research effort is targeted at the gut microbiome [1]. A number of diseases have been associated with alterations of individual microorganisms in the gut [1], and these associations are usually made through a statistical analysis commonly referred to differential abundance (DA) testing [2]. Differential abundance testing involves the use of statistical testing to determine if the relative abundances of certain microorganisms are significantly different between defined groups [3]. Numerous DA testing methods exist and include classical statistical tests (e.g. Kruskal-Wallis rank-sum test), methods originally developed to detect differential expression of gene transcripts in RNA-Seq data and adapted for microbiome analysis (e.g. DESeq2, edgeR), methods specifically designed for detecting differentially abundant microorganisms in microbiome data (e.g. ANCOM, metagenomeSeq), and methods designed to detect differentially abundant features (whether it be microorganisms or gene transcripts) in compositional data (e.g. ALDEx2). Differences in choice of DA method can contribute to inter-study variation in results, even between studies of the same disease, as most, if not all, methods will respond differently to microbiome data due to differences in their underlying characteristics. Multiple studies have previously assessed and compared the performance of popularly used DA testing methods, measuring their false positive rates (FPR), false discovery rates (FDR), sensitivities, and/or specificities using simulated data [2-5], with only one of these studies testing different methods on real data [4]. An example of how DA testing methods compare to one another when performed on real, complex disease gut microbiome datasets is still lacking in the literature. Also, not all methods included in previous comparison studies have been compared side by side as each study compared few to several methods at a time with slight differences in what methods were included in their assessments and comparisons.

Due to the lack of literature on how different DA methods behave and compare to one another when performed on real, complex disease gut microbiome datasets, DA testing was performed here using a variety of methods on two, large Parkinson disease (PD) – gut microbiome datasets in order to compare their results. Sixteen DA testing methodologies were found in the literature, and used to test for differentially abundant microbial genera (DA signatures) between PD and neurologically healthy controls in both datasets. Their results were then compared within and across datasets. Concordances between methods varied, although a subset of methods consistently resulted in higher concordances on average, especially with one another. These methods also detected DA signatures across datasets that seemed more robust to inter-methodological variation. Another subset of methods consistently produced the least concordant results on average with other methods, but some detected DA signatures in both datasets involving a subset of rare genera, all with higher relative abundances in PD. The vast majority of genera tested (∼80%) were detected as differentially abundant by at least one method in each dataset, but fewer signals were actually replicated in both datasets (46%), and even fewer were replicated by the majority (18%) or all (<1%) of the methods.

## Methods

### Subjects, metadata, gut microbiome

The original study was approved by institutional review boards at all participating institutions. Subjects, metadata, and gut microbiome data of datasets 1 and 2 have been previously described [6, 7]. We enrolled subjects and collected metadata and fecal samples from 212 PD and 136 neurologically healthy control subjects for dataset 1, and 323 PD and 184 neurologically healthy controls for dataset 2. Dataset 1 subjects were enrolled in Seattle, WA, Albany, NY, and Atlanta, GA, while all dataset 2 subjects were enrolled in Birmingham, AL. Methods for enrollment and collection of metadata and fecal samples were uniform across enrollment sites. PD was diagnosed according to the UK Brain Bank criteria by movement disorder specialists. Controls were self-reported free of neurological disease. Metadata were collected using questionnaires, or extracted from medical records. Stool samples were collected at home using DNA/RNA-free sterile swabs and mailed through U.S. postal service. All subjects provided written informed consent for their participation in the study.

DNA was extracted from stool samples using the automated MoBio PowerMag Soil DNA Isolation Kit (dataset 1) or manual MoBio PowerSoil DNA Isolation Kit (dataset 2). Hypervariable region 4 (V4) of the 16S rRNA gene was amplified with primers 515F-806R. Paired-end 150 bp (dataset 1) or paired-end 250 bp (dataset 2) sequencing was performed on V4 amplicons using Illumina MiSeq. Fifteen samples in dataset 1 resulted in low sequence count and were excluded.

Bioinformatic processing of sequences was performed separately for each dataset. Primers were trimmed from sequences using cutadapt v 1.16 [8]. DADA2 v 1.8 was used for quality trimming and filtering sequences, de-replicating sequences, inferring amplicon sequence variants (ASVs), merging of forward and reverse sequences, and detection and removal of chimeras [9]. Final ASV tables for dataset 1 and dataset 2 contained 6,844 unique ASVs for 201 PD and 132 controls samples and 12,198 unique ASVs for 323 PD and 184 control samples respectively. Taxonomy was assigned to ASVs using DADA2’s native implementation of the Ribosomal Database Project naïve Bayesian classifier with SILVA v 132 as reference and a bootstrap confidence of 80% [10]. Phylogenetic trees were constructed by first performing a multiple sequence alignment with DECIPHER v 2.8.1 [11], then building a phylogenetic tree with phangorn v 2.8.1 [12]. Phyloseq v 1.24.2 was used to create a phyloseq object for each dataset containing their respective ASV table, taxonomy classifications, phylogenetic trees, and subject metadata [13]. To agglomerate ASV level phyloseq objects to genus level, the tax_glom function in phyloseq was used without removal of unclassified genera. Total number of genera detected in dataset 1 was 445. Total number of genera detected in dataset 2 was 561.

### Differential abundance testing

We performed DA testing in dataset 1 and 2 separately using 16 DA methods. Method characteristics and parameters chosen for each method that differed from default can be found in Tables S1 and S2 respectively. Analyses using each method was performed as follows:

*Kruskal-Wallis rank sum test* [14]: Genera counts were transformed using total sum scaling, then unclassified genera, and genera present in < 10% of samples were excluded. The kruskal.test function from the stats R package was used to test for significant differences in genera relative abundances between PD and controls. *P* values were corrected for multiple testing using Benjamini-Hochberg (BH) FDR method implemented in the p.adjust function from stats package.

*Welch’s t-test with log transformation (log t-test)* [15]: Genera counts with a pseudo-count of 1 added were log transformed, then transformed using TSS. Unclassified genera, and genera present in < 10% of samples were excluded. The t.test function from the stats R package was used to test for significant differences in genera relative abundances between PD and controls. *P* values were corrected for multiple testing using BH FDR method from the p.adjust function.

*Negative binomial generalized linear model with and without zero-inflation (GLM NBZI):* Total sequence count was calculated for each sample. Unclassified genera, and genera present in <10% of samples were then excluded. Using raw counts, a negative-binomial generalized linear model with and without a zero-inflation component was fitted for each genus with the glmmTMB R package v 0.2.2.0 using log(total sequence count) as an offset variable, and PD vs control as the independent variable. Results were extracted from the model with the lowest Akaike information criterion. *P* values were calculated using the base summary function in R and corrected for multiple testing using BH FDR method implemented in the p.adjust function from stats package.

*Generalized linear model with centered-log-ratio transformed data (GLM CLR):* Genera counts with a pseudo-count of 1 added were centered-log-ratio (CLR) transformed, then unclassified genera, and genera present in < 10% of samples were excluded. A standard linear regression model using Gaussian distribution was fitted for each genus with the glm function from the R stats package with PD vs control as the independent variable. *P* values were calculated using the base summary function in R and corrected for multiple testing using BH FDR method implemented in the p.adjust function from stats package.

*Analysis of Composition of Microbes (ANCOM)* [16]: ANCOM was ran twice, once excluding unclassified genera, and genera present in < 10% of samples (ANCOM filtered), and again using all genera (ANCOM unfiltered), due to the drastic decrease in significant signals observed for ANCOM when filtering genera before analysis. Raw counts of genera were used as input to the ANCOM.main function from the ANCOM v 2 R code (downloaded from https://sites.google.com/site/siddharthamandal1985/research). PD vs control was specified as the main variable. The taxa-wise FDR option (multcorr=2) was chosen for the multiple testing correction method. An FDR significance threshold of 0.05 was chosen for calculation of *W* statistics. *W* statistics greater than or equal to 80% of the total number of genera tested were considered significant.

*metagenomeSeq zero-inflated Gaussian (fitZIG)* [17, 18]: Cumulative sum scaling (CSS) was applied to genera counts using the cumNorm function in metagenomeSeq R package v 1.22.0. Unclassified genera, and genera present in < 10% of samples were then excluded. A zero-inflated Gaussian model was fitted for each genus using function fitZig in metagenomeSeq. *P* values were corrected for multiple testing using BH FDR method implemented in the MRfulltable function in metagenomeSeq.

*metagenomeSeq fitFeatureModel* [17, 18]: CSS was applied to genera counts using cumNorm function in metagenomeSeq. Unclassified genera, and genera present in < 10% of samples were then excluded. A zero-inflated log-normal model was fitted for each genus using function fitFeatureModel in metagenomeSeq. *P* values were corrected for multiple testing using BH FDR method implemented in the MRfulltable function in metagenomeSeq.

*edgeR exactTest-TMM (edgeR TMM)* [19]: Using raw genera counts, normalization factors were calculated with the trimmed mean of M-values (TMM) method using the calcNormFactors function in edgeR R package v 3.22.5. Common and tagwise dispersions were then estimated using estimateCommonDisp and estimateTagwiseDisp functions in edgeR. Unclassified genera, and genera present in < 10% of samples were then excluded. Testing for differential relative abundance between PD and controls was performed using exactTest function in edgeR. *P* values were corrected for multiple testing using BH FDR method implemented in the topTags function in edgeR.

*edgeR exactTest-RLE (edgeR RLE)* [19]: Using genera counts with a pseudo-count of 1 added, normalization factors were calculated with the relative log expression (RLE) method using the calcNormFactors function in edgeR. The remaining steps were the same as edgeR TMM.

*DESeq2 nbinomWaldTest* [20]: Using raw genera counts, normalization factors were calculated using the function estimateSizeFactors in DESeq2 R package v 1.20.0 specifying type=”poscounts”. Unclassified genera, and genera present in < 10% of samples were then excluded. Testing for differential relative abundance between PD and controls was performed using the DESeq function in DESeq2 with default parameters. *P* values were corrected for multiple testing using BH FDR method implemented in the results function in DESeq2.

*limma-voom* [21]: Using raw genera counts, TMM values were calculated using the calcNormFactors function in edgeR. Log2-counts per million transformation and mean-variance trend estimation was performed using the voom function in limma R package v 3.36.5. Unclassified genera, and genera present in < 10% of samples were then excluded. Testing was performed by first fitting a linear model for each genus using function lmFit in limma, then testing for differential relative abundance between PD and controls using the eBayes function in limma. *P* values were corrected for multiple testing using BH FDR method implemented in the topTable function in limma.

*baySeq* [22]: Total sequence count was calculated for each sample. Unclassified genera, and genera present in < 10% of samples were then excluded. PD and control designations were used as the replicate structure. A list of two group structures was created where one group structure specified all subjects belonged to the same group, and the other specified PD and control groups. The replicate structure, list of group structures, and raw genera counts were combined into a countData object. Total sequence counts were manually supplied to the countData object. Priors were estimated from a negative binomial distribution using the function getPriors.NB in baySeq R package v 2.14.0, then likelihoods were estimated using function getLikelihoods in baySeq. FDR values were calculated using the topCounts function in baySeq.

*ALDEx2* [23]: Unclassified genera, and genera present in < 10% of samples were excluded. Raw genera counts were then used as input for the aldex function in ALDEx2 R package v 1.12.0 specifying 1000 Monte Carlo samples. Both Wilcoxon (ALDEx2 Wilcoxon) and t-test (ALDEx2 t-test) were used for testing differences in genera relative abundances between PD and controls. *P* values were corrected for multiple testing using BH FDR method implemented in the aldex function.

*SAMseq* [24]: The SAM method for normalization of sequence counts (Anscombe transformation, then dividing by the square root of sequencing depth) was applied to genus counts using the samr.norm.data function in the samr R package v 3.0. Normalized values were rounded to the nearest integer. Unclassified genera, and genera present in < 10% of samples were excluded. Normalized genera counts were then used as input for the SAMseq function in the samr R package specifying “Two class unpaired” as the response type and the fdr.output as “1” in order to get full result list. FDR q-values were extracted from “siggenes.table” in the SAMseq output.

*Linear discriminant analysis Effect Size (LEfSe)* [25]: Genera counts were transformed using TSS, then unclassified genera, and genera present in < 10% of samples were excluded. Sample IDs, case/control class designations, and genera relative abundances were exported from R and used as input for LEfSe v 1.0.8.post1 (downloaded using LEfSe bioconda recipe https://bioconda.github.io/recipes/lefse/README.html). Only genus level taxonomy designations were included in the LEfSe input. The LEfSe input was formatted using the lefse-format_input.py script specifying the normalization value to be “1E6”. LEfSe analysis was then ran on the formatted data using the run_lefse.py script with default parameters. Since LEfSe only outputs uncorrected *P* values for features that it finds significant, LEfSe analysis was ran again, but this time specifying parameters that would output all *P* values. The full range of LEfSe *P* values were multiple testing corrected using BH FDR method implemented in the p.adjust function from stats package. These corrected *P* values were substituted for the uncorrected *P* values outputted by the default LEfSe run.

After exclusion of unclassified genera and genera found in <10% of subjects, 109 and 163 genera remained for DA testing in dataset 1 and 2 respectively, with 106 genera in common between both datasets. Unless otherwise mentioned above, significance was set at FDR < 0.05. Significant differences in a genus’ relative abundance between PD and control groups will be referred to as a “DA signature” for the purpose of this manuscript. For a particular method, a DA signature was considered replicated in both datasets if it reached multiple testing corrected significance in both datasets for that method.

### Concordance of DA signatures across differential abundance methods

To measure similarity in DA signature calls between methods, pairwise concordances were calculated for each method. To calculate pairwise concordances between methods for each dataset individually, a binary genus by method matrix was first created with values denoting which methods did (1), or did not (0), detect a DA signature between a genus and PD. Then, for each pair of methods in turn, concordances were calculated by summing the number of values that were the same between both methods (both having 1 or 0 for a particular genus) and dividing by the total number of genera tested. Concordances for ANCOM unfiltered were calculated after removing results for genera not tested by the other methods (those found in <10% of samples, and unclassified at genus level). To calculate pairwise concordances between methods when taking replication of DA signatures across datasets into account, a genus by method matrix was first created with values denoting which methods detected a DA signature in no datasets (0), dataset 1 only (1), dataset 2 only (2), both datasets with opposite effect directions (3), or both datasets with same effect direction (4). The mean relative abundance ratio of PD to control subjects was used to determine effect direction. Only tested genera that were in common between both datasets were included in the matrix (106 genera). Then, for each pair of methods in turn, concordances were calculated by summing the number of values that were the same between both methods (both having a 0, 1, 2, 3, or 4) and dividing by the total number of genera tested and in common between datasets. Matrices of pairwise concordances were visualized as a heatmap using the ggcorrplot function from the ggcorrplot R package v 0.1.3. Cells were colored by a blue (lower concordance) to white (around the mean pairwise concordance of each heatmap) to orange (higher concordance) color gradient. Methods were ordered from lowest (bottom, left) to highest (top, right) mean pairwise concordance. Mean pairwise concordance for each method was calculated in order to see how similar a DA method’s calls were compared to all other methods on average. Methods with a mean pairwise concordance lower than the mean pairwise concordance of all methods in a particular dataset or combined results were considered to be part of a “lower concordant” group, while methods with a higher than average mean pairwise concordance was considered to be part of a “higher concordant” group.

To determine if any groups of DA signatures were being agreed (or disagreed) upon by all or a subset of methods, hierarchical clustering was performed to group DA signatures based on similarities in DA signature calls between methods. A genus by method matrix denoting which methods detected a DA signature between a genus and PD in no datasets (0), dataset 1 only (1), dataset 2 only (2), both datasets with opposite effect directions (3), or both datasets with same effect direction (4) was first created as stated above, then hierarchical clustering was performed and visualized in a heatmap using the heatmap.3 function (downloaded from https://raw.githubusercontent.com/obigriffith/biostar-tutorials/master/Heatmaps/heatmap.3.R on 10/21/2019) with the default distance function, but specifying the hierarchical clustering function (hclustfun) to be diana from the cluster v 2.1.0 R package. DIANA (DIvisive ANAlysis) performs a divisive hierarchical clustering algorithm [26], which, in this situation, attempts to group DA signatures based on the similarities in DA signatures being detected in either no datasets, dataset 1 only, dataset 2 only, both datasets with opposite effect directions, or both datasets with same effect direction between methods. The PD to control mean relative abundance ratios (MRAR) and control mean relative abundances (MRA) for each genus were also plotted next to the heatmap. MRARs were given a color gradient from red (lowest MRAR) to white (MRAR ∼ 1) to blue (highest MRAR). Control MRAs were given a color gradient from white (lowest MRA) to dark green (highest MRA). Hierarchical clustering was also performed for methods in an effort to arrange and group them based on their result similarities.

### Testing for differences between datasets and groups

To test for differences between datasets and groups detected by hierarchical clustering, t-tests were performed and results recorded in the Supplementary Information section of the Supplementary Data.

## Results

### Method characteristics

Methods included in the present study span the fields of traditional statistics, RNA-Seq analysis, and microbiome analysis, and have varying underlying characteristics. A summary of method characteristics for the 16 DA methods can be found in Table S1. The majority of methods included here utilized parametric statistical tests (assumes the data has some form of underlying distribution). Of these, the most commonly assumed data distribution was the negative binomial distribution (DESeq2, baySeq, edgeR RLE, edgeR TMM, GLM NBZI). No data transformations were performed for negative binomial methods, or metagenomeSeq methods, to try and bring the data to normality as non-normality of data is taken into account in their statistical models. The remaining parametric methods (ALDEx2 t-test, log t-test, limma-voom, GLM CLR) all used statistical tests that assumed a Gaussian distribution of the data, therefore, transformations were needed before analysis, which here included a log transform of some kind. Five methods (ALDEx2 Wilcoxon, ANCOM, Kruskal-Wallis, SAMseq, LEfSe) were considered non-parametric (assumes no underlying distribution of data) as they used statistical tests that transformed data to ranks. Methods also differed in what techniques were used to account for varying sequence depth between samples. Four of the five negative binomial methods (DESeq2, baySeq, edgeR RLE, edgeR TMM) calculated scaling factors for each sample to account for uneven sequence count. Cumulative sum scaling was used for both metagenomeSeq methods. Total sum scaling (also referred to as relative abundance) was performed for three methods that did not have a built in technique to account for varying sequence depth (Kruskal-Wallis, log t-test, LEfSe) as it is a widely used normalization technique and recommended as the normalization technique of choice by LEfSe authors [25]. Log ratio transformations were used for ALDEx2 methods, GLM CLR, and ANCOM, which, in addition to normalizing for inter-sample variation in total sequence count, takes the compositionality of the data into account. The remaining 3 methods (limma-voom, GLM NBZI, SAMseq) did not share normalization techniques with any other methods. To account for varying sequencing depth, limma-voom utilized a log2-counts per million transformation, the log of the sequencing depth was used as an offset variable in the model for GLM NBZI, and SAMseq utilized its own unique transformation which included an Anscombe transformation followed by dividing by square root of the sequencing depth. The majority of methods (14 out of 16) have been previously included in method comparison studies [2-5], and all seemed to keep FPR and/or FDR under 0.2 when performed on simulated data except for fitZIG, edgeR, and SAMseq, which were show to reach upwards of 0.9 (Table S1).

### Concordance of DA signatures between methods

On average, DA signature calls between methods varied in concordance, with pairwise concordances between methods ranging from 46%-99% similarity with the mean pairwise concordance being 76% per dataset (Fig. 1, Table S3). For both datasets, results from baySeq, GLM NBZI, fitZIG, limma-voom, edgeR, and SAMseq had the lowest pairwise concordances on average with other methods (61%-76%). Results from Kruskal-Wallis, log t-test, GLM CLR, fitFeatureModel, ALDEx2, DESeq2 and LEfSe had the highest pairwise concordances on average with other methods (78%-82%) especially among one another (88%-89%). When incorporating information on what DA signatures were replicated across datasets by a given method, the mean pairwise concordance between all methods decreased compared to individual datasets (62%). As seen in individual datasets, baySeq, GLM NBZI, fitZIG, limma-voom, edgeR, and SAMseq had the least similar calls on average with other methods (46%-59%), and Kruskal-Wallis, log t-test, GLM CLR, fitFeatureModel, ALDEx2, DESeq2 and LEfSe had the most similar calls on average with other methods (62%-70%), and among each other (77%). Although the two datasets analyzed here have been previously reported to have significant heterogeneity in microbiome composition [7], no significant difference between datasets was found for average pairwise concordance (*t*(30)=0.3, *P*=0.78), but there was a significant decrease in average pairwise concordance when accounting for replication of signals in both datasets (76% per dataset vs 62% with replication; *P* < 1E-6). This indicates that concordances between methods will inevitably fall when requiring DA signatures to replicate across two datasets. This may be attributed to variation in underlying technical and population characteristics between datasets, heterogeneity in microbiome compositions between different populations, and/or false positives detected in a single dataset that are irreproducible in additional datasets.

**Fig 1.**
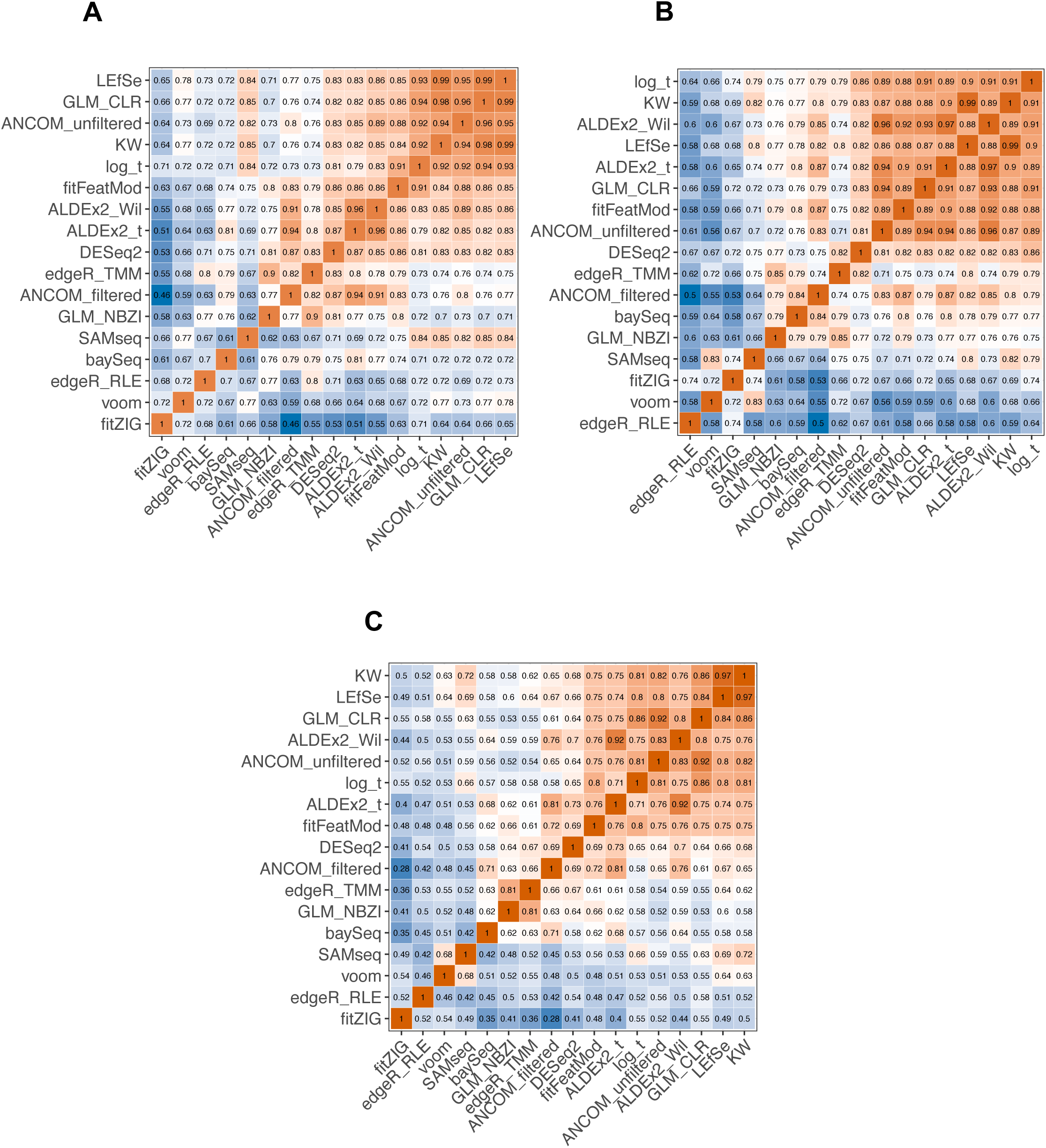
Pairwise concordances between methods. Pairwise concordances were calculated between method DA signature calls for (A) 109 genera in dataset 1, (B) 163 genera in dataset 2, and (C) results when accounting for replication across datasets for 106 genera in common between both datasets. Values in heatmap cells correspond to the concordance between two methods. Concordances were calculated by summing the number of differential abundance signatures that were called the same between two methods (i.e both calling a differential abundance signature significant or not significant for (A) and (B), or significant in either no datasets, dataset 1 only, dataset 2 only, both datasets with opposite effect directions, or both datasets with same effect direction for (C)) and dividing by the total number of genera tested. Concordances for ANCOM_unfiltered were calculated after removing results for genera not tested by the other methods (those found in <10% of samples, and unclassified at genus level). Cells are colored by a blue (lower concordance) to white (around the mean pairwise concordance of each heatmap) to orange (higher concordance) color gradient. The mean pairwise concordance was 0.76 for (A) and (B), and 0.62 for (C). Methods are ordered from lowest (bottom, left) to highest (top, right) mean pairwise concordance. KW: Kruskal-Wallis; GLM_CLR: generalized linear model with centered log ratio transformation; ALDEx2_Wil: ALDEx2 with Wilcoxon rank-sum test; log_t: Welch’s t-test with log transformation; ALDEx2_t: ALDEx2 with t-test; fitFeatMod: fitFeatureModel from metagenomeSeq; edgeR_TMM: edgeR exact test with trimmed mean of M-values; GLM_NBZI: generalized linear model assuming negative binomial distribution with, or without, zero-inflation; edgeR_RLE: edgeR exact test with relative log expression.

### Variable effect of input data filtering for ANCOM

When performing analyses with ANCOM after excluding unclassified genera and genera found in <10% of subjects (referred to as ANCOM filtered here), we observed that very few DA signatures were being detected. DA signature calls for ANCOM filtered resulted in below average pairwise concordances, placing it in the lower concordant group of methods mentioned above for both datasets and when replication of signals were taken into account (Fig. 1). ANCOM shares similar characteristics with methods included in the higher concordant group of methods (non-parametric like Kruskal-Wallis, LEfSe, ALDEx2 Wilcoxon, and uses a log ratio transformation like GLM CLR and ALDEx2), therefore, it was unexpected when it fell into the lesser group. To investigate the effect of input data filtering on ANCOM results, ANCOM was performed again using all genera in the analysis (referred to as ANCOM unfiltered here). In both datasets, comparing results between ANCOM filtered and ANCOM unfiltered showed a reduction in the detection of DA signatures when filtering genera before analysis (82% reduction in dataset 1 and 79% reduction in dataset 2, Table S4). We then investigated the effect of filtering for two other standard statistical methods (GLM CLR and Kruskal-Wallis) by performing each on filtered and unfiltered data. Both GLM CLR and Kruskal-Wallis detected more DA signatures when data was filtered before analysis compared to when all genera were included in analysis (Table S4). The changes in detected DA signatures for GLM CLR and Kruskal-Wallis were less severe than ANCOM’s with GLM CLR and Kruskal-Wallis detecting ∼29% and ∼33% more DA signatures with filtered data in both datasets respectively. Pairwise concordances between ANCOM unfiltered and other methods were also higher on average than those seen with ANCOM filtered, placing ANCOM unfiltered among the group of methods with higher mean pairwise concordances (Fig. 1, Table S3). Because DA signature calls made by ANCOM unfiltered were more similar to other methods on average, we used results from ANCOM unfiltered in place of ANCOM filtered for further comparisons between method results.

### Comparison of method results across datasets

Datasets differed in size and had significant heterogeneity in microbiome composition [7], therefore, we compared method calls across datasets to see if there were any significant dataset differences in the number or proportion of DA signatures being detected by the 16 methods. Within each dataset, ∼80% of genera were detected as differentially abundant by at least one method, 26%-27% of genera were detected as differentially abundant by at least half of the methods, and 3%-4% by all 16 methods (Tables S5 and S6). No significant difference was found between datasets for the average number of methods that detected a particular DA signature (*t*(235)=-0.4, *P*=0.67). When comparing results across datasets for replications, 46% (49/106) and 18% (19/106) of genera tested and in common between datasets were detected as differentially abundant by at least one method or half of the methods respectively (Table S7). Only one genus (*Agathobacter*) was detected by all 16 methods in both datasets.

The number of DA signatures detected by a particular method in each dataset ranged from a small minority to over 50% of tested genera. For dataset 1, the maximum number of DA signatures detected by a particular method was 64 (fitZIG, encompassing 59% of tested genera), the minimum detected was 11 (ALDEx2 t-test, encompassing 11% of tested genera), and the mean per method was 31±14 (encompassing on average 28% of tested genera) (Table S5). For dataset 2, the maximum number of DA signatures detected by a particular method was 90 (edgeR RLE, encompassing 55% of tested genera), the minimum detected was 30 (ALDEx2 t-test, encompassing 18% of tested genera), and the mean per method was 50±21 (encompassing on average 31% of tested genera) (Table S6). Methods on average detected a significantly higher number of DA signatures in the larger, deeper sequenced dataset 2 (*t*(26)=-2.9, *P*=7E-3), but no significant difference was found between datasets when the DA signature count for each method was normalized by the number of genera tested in analysis (*t*(30)=-0.4, *P*=0.72). When taking replication of signals into account, the maximum number of DA signatures replicated by a particular method was 37 (fitZIG, encompassing 35% of tested genera in common between datasets), the minimum detected was 8 (baySeq, encompassing 8% of tested genera in common between datasets), and the mean per method was 20±9 (encompassing on average 19% of tested genera in common between datasets) (Table S7).

Overall, despite differences in the size of datasets and heterogeneity in microbiome composition, we observed no significant differences between datasets in the proportion of genera being detected as differentially abundant on average. The number of detected DA signatures per method was significantly increased in dataset 2, but that is to be expected since it is a larger, more powered dataset both in sample size and number of genera detected. As expected, requiring replication across datasets decreased the number DA signatures being detected per method, which might be a reflection of weeding out potentially false positive, or heterogeneous signals.

### Hierarchical clustering of genera and methods based on similarities in DA signatures

To observe what groups of DA signatures either all, or subsets, of methods were agreeing, or disagreeing, upon, we performed hierarchical clustering of DA signatures and methods based on similarities in DA signature calls and replications (Fig. 2). Hierarchical clustering grouped DA signatures into three groups. The first group encompassed 26 genera (25% of genera tested and in common between datasets) that were more likely to be detected as differentially abundant across methods in both datasets (10±4 methods on average; Fig. 2, group 1, Table S7). This group included genera that were found to be both enriched or depleted in PD, and had a wide range of control MRAs and effect sizes ranging from highly prevalent genera with moderate effect sizes (e.g. *Agathobacter,* control MRA = 0.02-0.04, absolute MRAR = 1.8 – 1.9) to rarer genera with larger effect sizes (e.g. *Prevotella*, control MRA = 6E-4 – 2E-3, absolute MRAR = 2.6 – 4.4). The second group included 67 genera (63% of genera tested and in common between datasets) that were detected as differentially abundant in both datasets by little to no methods (<1 methods on average; Fig. 2, group 2, Table S7). On average, group 2 included more prevalent genera (mean control MRA = 0.01) with smaller effect sizes (mean absolute MRAR = 1.3 – 1.5) when compared to the first group (mean control MRA = 5E-3 – 6E-3, mean absolute MRAR = 2 – 2.6). The third group included 13 genera (12% of genera tested and in common between datasets), all found to be enriched in PD, who were more likely to be detected as differentially abundant by a specific subset of methods (4±2 methods on average; Fig. 2, group 3, Table S7). Group 3 was interesting as it contained only genera that were enriched in PD, had higher effect sizes than the previous two groups (mean absolute MRAR = 3.4 – 3.8), was made up of genera with very low control MRAs (mean control MRA = 7E-4), and were detected in both datasets by only a subset of methods that were included in the lower concordant group of methods previously mentioned above.

**Fig. 2.**
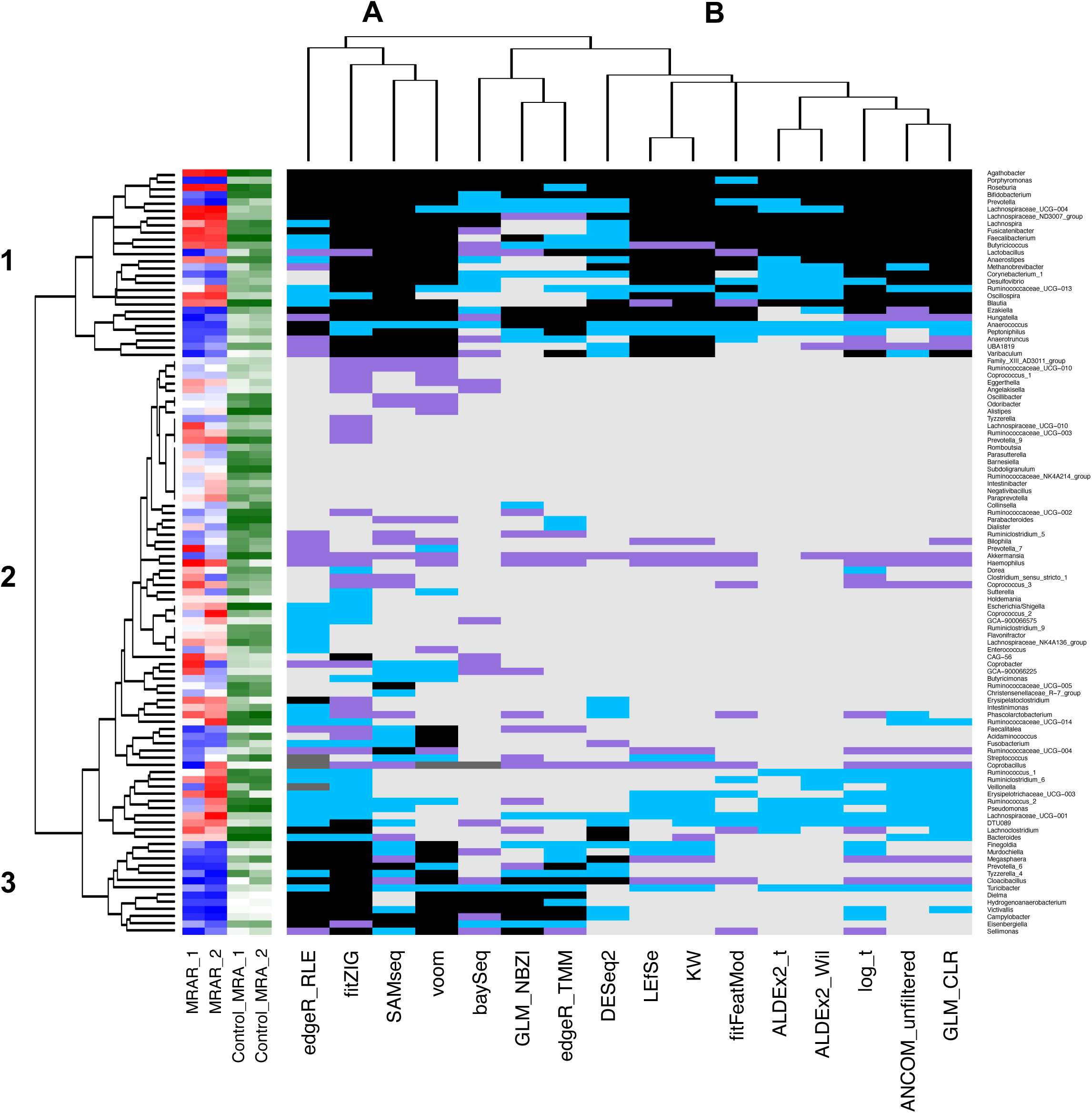
Hierarchical clustering of genera and methods based on similarity in detected differential abundance signatures. Hierarchical clustering was performed to group genera (rows) and methods (columns) based on similarities in detected and replicated differential abundance signatures and was visualized via heatmap. Three groups of genera were revealed by hierarchical clustering: (1) genera more likely to be called differentially abundant by the majority of methods in both datasets, (2) genera who were called differentially abundant in both datasets by little to no methods, and (3) genera enriched in PD with low relative abundances who were called differentially abundant in both datasets by a subset of methods. Only results for genera tested and in common between datasets (106 genera) were included in hierarchical clustering and heatmap. Cells correspond to a differential abundance signature that was detected in no datasets (value=0, color=light gray), dataset 1 only (value=1, color=purple), dataset 2 only (value=2, color=light blue), both datasets with opposite effect directions (value=3, color=dark grey), or both datasets with same effect directions (value=4, color=black). Mean relative abundance ratios for genera in dataset 1 (MRAR_1) and dataset 2 (MRAR_2) were plotted next to the heatmap, and given a color gradient from red (lowest MRAR) to white (MRAR ∼ 1) to blue (highest MRAR). Control mean relative abundances for dataset 1 (Control_MRA_1) and dataset 2 (Control_MRA_2) were also plotted next to the heatmap, and given a color gradient from white (lowest MRA) to dark green (highest MRA). KW: Kruskal-Wallis; GLM_CLR: generalized linear model with centered log ratio transformation; ALDEx2_Wil: ALDEx2 with Wilcoxon rank-sum test; log_t: Welch’s t-test with log transformation; ALDEx2_t: ALDEx2 with t-test; fitFeatMod: fitFeatureModel from metagenomeSeq; edgeR_TMM: edgeR exact test with trimmed mean of M-values; GLM_NBZI: generalized linear model assuming negative binomial distribution with, or without, zero-inflation; edgeR_RLE: edgeR exact test with relative log expression.

Hierarchical clustering of methods revealed two groupings of methods. The first group contained more than half of the lesser concordant methods (edgeR RLE, fitZIG, SAMseq, limma-voom; Fig. 2, group A, Table S7) that, together, detected the majority of genera included in group 3 as differentially abundant across datasets. The second group contained the remainder of methods (Fig. 2, group B, Table S7). Group A detected significantly more genera on average as differentially abundant in one or more datasets compared to group B (69±9 vs 40±6, *t*(4)=6.2, *P*=3E-3) including DA signatures encompassed in group 2 that had the least consensus among methods (31±8 vs 11±4, *t*(3)=4.8, *P*=0.02). No other groupings of methods seemed obvious from the hierarchical clustering, but of note, the next major split in the dendrogram for group B was between the remaining lesser concordant methods not included in group A (baySeq, GLM NBZI, edgeR TMM) that also detected and replicated DA signatures in group 3, and the higher concordant methods (DESeq2, LEfSe, Kruskal-Wallis, fitFeatureModel, ALDEx2, log t-test, ANCOM unfiltered, GLM CLR).

## Discussion

In summary, 16 differential abundance testing methods were found in the literature and used to detect differentially abundant genera in PD patients versus healthy controls in two large PD-gut microbiome datasets. Methods spanned multiple fields and had both common and unique characteristics when compared to one another. Pairwise concordances between method calls varied overall. Certain methods consistently resulted in higher mean pairwise concordances (e.g. Kruskal-Wallis, ALDEx2, GLM CLR), while others consistently resulted in lower mean pairwise concordances (e.g. edgeR, fitZIG). Differential abundance signatures were detected by all methods and the number of DA signatures detected by each method ranged from a small subset of genera to over half of the genera tested. Overall, ∼80% of genera tested were associated with PD by at least one method in each dataset. Requiring associations to replicate across datasets reduced significant signals by almost half. Further requirement of signals to be replicated by the majority of methods (≥8) yielded 19 significant associations. Only one genus (*Agathobacter*) was replicated by all methods. Hierarchical clustering revealed three groups of DA signatures that were (1) replicated by the majority of methods and included genera previously associated with PD, (2) replicated by few or no methods, and (3) replicated by a subset of methods and included rarer genera, all enriched in PD. Although datasets were heterogeneous in microbiome composition, we observed no significant differences between datasets in average concordances and the proportion of genera being detected as differentially abundant on average.

The variation between method results reported here aligns with the variation between differential abundance testing method performances previously reported in method comparison studies [3-5]. Performance of methods could not be assessed here, as analyses were conducted on real datasets where the true answers are unknown, but it was observed that certain methods consistently resulted in above average pairwise concordances, and even higher pairwise concordances among each other, while other methods consistently resulted in lower than average pairwise concordances. Interestingly, methods found here to have above average pairwise concordance were also previously reported to have lower FPR and/or FDR (less than 0.1; e.g. Kruskal-Wallis, log t-test, ALDEx2, fitFeatureModel, ANCOM, DESeq2). This, along with observations made in the current study, might help explain why they resulted in such high concordances with each other. Methods with lower FPR/FDR have been previously reported to be conservative in their performance [3], detecting less microorganisms as differentially abundant compared to higher FPR/FDR methods (which we also observed in our data). In this study, these methods seemed to detect more DA signatures across datasets that were robust to inter-methodological variation (i.e. genera shown in Fig. 2, group 1 that were more likely to be replicated by the majority of methods on average). Taken together, we can extrapolate that methods with lower FPR and/or FDR might be more likely to converge on the same microorganisms because they are detecting less microorganisms as differentially abundant overall, and the signatures they do detect might be true signals that are more robust to methodological variation and reproducible across datasets. Based on results of this study, these methods seem to be good candidates for detection of DA signatures that will be reproducible across DA testing methods and different datasets. This comes with the caveat that some true associations might be missed as these methods seem to be detecting more “high confidence” hits, and therefore, might be conservative as previously stated.

A surprising finding from this study was the variable effect of input data filtering prior to testing with ANCOM. Usually, filtering of taxonomic data before analysis would increase the number of DA signatures detected by a method, mostly due to the decreased burden of multiple testing correction at the FDR calculation step. However with ANCOM, filtering of input data before analysis greatly decreased the number of detected DA signatures. This might be due to the different statistics used by ANCOM compared to the standard FDR q-value. To determine significance, ANCOM calculates a *W* statistic, which is the number of times the log ratio of a microorganism with every other microorganism being tested was detected to be significantly different across groups (in this case PD vs control) [16]. Because *W* statistics are based on pairwise comparisons between all microorganisms being tested, they will automatically decrease overall if less microorganisms are included in the analysis, and the threshold range for significant *W* statistics will also decrease. In addition, if low prevalent microorganisms are being removed, this will not only decrease the *W* statistics overall, but now *W* statistic calculation might become more conservative since higher prevalent, potentially more stable microorganisms have been selected for, the ratios of which might not differ enough to be detected as significant at a particular *W* statistic threshold. This could potentially make the analysis overly conservative (as seen in this study), therefore, it might be beneficial to perform ANCOM using the full surveyed microbiome, removing none, or only the very rare microorganisms, before analysis.

A finding from this study that helped illustrate the behavior of the methods on the data analyzed here was the detection of two groups of genera that were converged upon by either the majority or subset of methods. Hierarchical clustering of genera based on similarity in method results showed one group of genera that were more likely to be detected as differentially abundant in both datasets by the majority of methods on average (Fig. 2, group 1). Theoretically, this group might be looked at as the “high confidence” group, as methods from across the spectrum tended to detect the genera in this group as differentially abundant in both datasets. This group included genera previously associated with PD such as *Bifidobacterium*, *Lactobacillus*, and short-chain fatty-acid producing bacteria *Faecalibacterium*, *Roseburia*, *Blautia*, and other members of the *Ruminococcaceae* and *Lachnospiraceae* family [27, 28]. It also included members of a correlated poly-microbial group of genera found previously to be increased in PD (*Porphyromonas*, *Corynebacterium_1*, *Prevotella*, *Ezakiella*, *Peptoniphilus*, *Anaerococcus*, *Varibaculum*) [7]. Hierarchical clustering also revealed a second group of genera that were only detected as differentially abundant in both datasets by a subset of methods (fitZIG, edgeR, limma-voom, baySeq, SAMseq, GLM NBZI, DESeq2; Fig. 2, group 3). This group was intriguing because it only contained genera enriched in PD that had low control MRAs, and higher effect sizes on average compared to groups 1 and 2. Without the use of these methods, this group of genera might have been missed, arguing that, although some of these methods were previously reported to have higher FPR/FDR [3-5], they might detect associations that may be missed by more conservative methods. It seems these methods should be used with caution, however, as they tended to detect and replicate a large portion of genera as differentially abundant, even those that were included in the DA signature sparse group 2. As done here, the use of a second, confirmatory dataset might assist in helping curve the number of false positives these methods could potentially detect, and reveal which associations are reproducible. However, the caveat is that, because a number of these methods detected a large portion of the tested genera as differentially abundant in both datasets (e.g. fitZIG and edgeR), the chance of replication across datasets is automatically increased, therefore, false positives may still be possible even if requiring DA signatures to replicate across datasets.

One of the biggest limitations of this study is the lack of ability to test the actual performance of these methods, as no simulations were performed and only real data was used, so the true answers are unknown. Luckily, the majority of the methods studied here have been previously subjected to comparison studies on simulated data, and results from those studies could be used to inform the discussion of the results from this study [2-5]. Unfortunately, not all methods implemented in this study had previously reported performance metrics (i.e. LEfSe, GLM CLR, GLM NBZI), therefore, it is difficult to speculate reasons for why they resulted in higher or lower concordances with other methods. A methodological limitation of this study is the choice of parameters used for each method. Each method contains multiple functions with multiple parameters, and it was beyond the scope of this study to try different combinations of parameters to fully optimize each method for the data being analyzed. We attempted to make parameter choices based on what was default for the method and/or what was recommended for the method by the method authors especially in the context of microbiome data analysis, but this process is inherently biased as we did not attempt to try every combination of parameter choice possible for each method. Because this study was performed on two datasets collected for the study of a specific disease in a specific aged population with data derived from a specific host source (stool), the results reported here might not translate to all types of microbiome datasets. Both datasets analyzed here were also created using uniform methodology, therefore, results may also change if attempting to analyze two datasets created using different methodologies. The genera detected in this study as differentially abundant between PD and control subjects should be interpreted with caution as the goal of this study was to compare the results of different DA methods when performed on microbiome datasets of real, complex disease, and did not take into account any confounding variables that might drive false signals. Because no confounders were taken into account during the analyses of this study (e.g. PD medications, constipation, age, sex, diet), we cannot report differentially abundant genera detected in this study as truly “associated” with PD as the proper steps to guard against false positives due to underlying study population characteristics were not taken. Still, majority of genera listed as part of group 1 have been previously reported as associated with PD [7], and genera listed as part of group 3 provide interesting findings that will hopefully be elucidated further in future investigations.

## Conclusion

In conclusion, we performed 16 differential abundance testing methods in two large PD-gut microbiome datasets and compared their results. We found results varied between methods, but we found some methods resulted in higher pairwise concordances, while others resulted in lower concordances. Filtering of genera before analysis with ANCOM drastically reduces the number of DA signatures detected, most likely due to the way *W* statistics are calculated and used for significance. This suggests it might be more advantageous to supply ANCOM with unfiltered abundances for higher taxonomic levels such as genus, and only filter out very rare features (that are most likely just noise) for lower taxonomic levels. The majority of methods converged on a group of DA signatures that seemed more robust to inter-methodological differences, while a subset of methods converged on a smaller, second group of DA signatures involving genera enriched in PD with low relative abundances. This study helps to fill a void in the literature on how different DA methods behave when performed on real, complex disease oriented gut microbiome datasets, and hopefully it will help inform future studies looking to perform these types of analyses, especially those working with gut microbiome data derived for studying Parkinson disease.

## List of Abbreviations

DA: differential abundance PD – Parkinson disease
GLM: generalized linear model FPR – false positive rate
FDR: false discovery rate TSS – total sum scaling BH – Benjamini-Hochberg
NBZI: negative binomial zero-inflated CLR – centered-log-ratio
CSS: cumulative sum scaling TMM – trimmed mean of M-values RLE – relative log expression
MRAR: mean relative abundance ratio MRA – mean relative abundance

## Availability of data and materials

Individual-level raw sequences and basic metadata are publicly available at NCBI Sequence Read Archive (SRA) BioProject ID PRJNA601994. Scripts and phyloseq used in this study can be found at the following link: https://github.com/zwallen/Wallen_DAMethodCompare_2021. Full results for each method can be found in Tables S5 and S6.

## Author contributions

Z.D.W contributed to the conception and design of the work, the analysis and interpretation of the data, and drafting of the work.

## Ethics approval and consent to participate

The original study was approved by institutional review boards at all participating institutions. All subjects provided written informed consent for their participation in the study.

## Consent for publication

Not applicable.

## Acknowledgements

I would like to acknowledge Dr. Haydeh Payami as this work was performed under her mentorship and thank her for her guidance and helpful assistance through execution of the work and preparation of the manuscript.

